# σ^54^ (σ^L^) plays a central role in carbon metabolism in the industrially relevant *Clostridium beijerinckii*

**DOI:** 10.1101/505099

**Authors:** Rémi Hocq, Maxime Bouilloux-Lafont, Nicolas Lopes Ferreira, François Wasels

**Affiliations:** IFP Energies Nouvelles, 1 et 4 avenue de Bois-Préau, 92852 Rueil-Malmaison, France

**Keywords:** *Clostridium beijerinckii*, sigma factor, metabolism, metabolic engineering, Isopropanol-Butanol fermentation

## Abstract

Microbial production of butanol and isopropanol, two high value-added chemicals, is naturally occurring in the solventogenic *Clostridium beijerinckii* DSM 6423. Despite its ancient discovery, the precise mechanisms controlling alcohol synthesis in this microorganism are poorly understood. In this work, an allyl alcohol tolerant strain obtained by random mutagenesis was characterized. This strain, designated as the AA mutant, shows a dominant production of acids, a severely diminished butanol synthesis capacity, and produces acetone instead of isopropanol. Interestingly, this solvent-deficient strain was also found to have a limited consumption of two carbohydrates and to be still able to form spores, highlighting its particular phenotype. Sequencing of the AA mutant revealed point mutations in several genes including CIBE_0767 (*sigL*), which encodes the σ^54^ sigma factor. Complementation with the wild-type *sigL* gene fully restored solvent production and sugar assimilation, demonstrating that σ^54^ plays a central role in regulating these pathways in *C. beijerinckii* DSM 6423. Genomic comparison with other strains further revealed that these functions are probably conserved among the *C. beijerinckii* strains. The importance of σ^54^ in *C. beijerinckii* was further assessed by the characterization of a *sigL* deletion mutant of the model strain NCIMB 8052 obtained with a CRISPR/Cas9 tool. The resulting mutant exhibited phenotypic traits similar to the AA strain, and was subsequently complemented with the *sigL* gene from either the wild type or the AA strains. The results of this experiment confirmed the crucial role of σ^54^ in the regulation of both solventogenesis and sugar consumption pathways in *C. beijerinckii*.

**Importance:** *Clostridium beijerinckii* shows a significant potential for producing valuable biochemicals and biofuels. One of the major hurdles impeding its widespread usage is its low endogenous production of alcohols, which could be alleviated by metabolic engineering approaches. Despite its former long-time use in the industrial acetone-butanol-ethanol process, the molecular mechanisms controlling solventogenesis in the *Clostridium* genus still remain elusive, preventing genetic engineering approaches for strain enhancement. In this context, our study provides novel insights into the crucial role of the σ^54^ transcriptional factor in solvent synthesis regulation in two *C. beijerinckii* strains. Furthermore, we show that this sigma factor also controls sugar consumption and is therefore a key controller of carbon metabolism in this species.

## Introduction

In the context of worldwide energy transition, research for alternatives to fossil fuels has become a priority. In particular, the replacement of petrochemistry by a low carbon emission industry has been a major challenge as our global consumption of petrochemicals keeps on increasing (1). The valorization of plant biomass to synthesize ethanol by microbial fermentation has already been pioneered for biofuel production (2) and could therefore be applied to bio-based chemistry (3).

A few strains from the *Clostridium* genus are naturally able to produce isopropanol and butanol (4, 5), two compounds that could be used as biochemical and biofuel, respectively. However, those organisms are not producing these metabolites in quantities compatible with an economically viable industrial process (6). However, with the increasing availability of efficient genetic tools in *Clostridia* (7), metabolic engineering approaches could be undertaken to enhance solvent productivity. *Clostridium beijerinckii* DSM 6423 (NRRL B-593) is the only natural isopropanol-butanol producing strain whose genome and transcriptome have been investigated (8). It may therefore be the best candidate for genetic engineering, although its particular physiology is still poorly understood. For this purpose, gaining additional knowledge on metabolism regulation in this strain would greatly benefit future optimization efforts. In particular, identifying the molecular effectors controlling solvent production may provide valuable insights to define adequate genetic engineering strategies.

However, the limited genetic toolbox available for this particular strain has been a severe limitation. To circumvent this problem, Máté de Gerando and coworkers performed random mutagenesis coupled with genome shuffling to increase isopropanol productivity by selecting isopropanol-tolerant strains (9). In this work, random mutagenesis followed by allyl alcohol selection also generated an interesting mutant, further referred to as AA mutant. This strain mainly produces acids, shows no isopropanol production and a strongly attenuated butanol synthesis capacity. These results are consistent with those obtained in *Clostridium acetobutylicum* DSM 1792, in which mutants obtained in the presence of allyl alcohol - precursor of the highly toxic acrolein molecule in the reaction catalyzed by alcohol dehydrogenases - permitted the selection of butanol-deficient strains (10). Nevertheless, in both cases the key mutated genes causing these phenotypes have not been clearly identified.

In this study, we demonstrate that the solvent production deficiency in *C. beijerinckii* DSM 6423 AA is due to a point mutation in the CIBE_0767 gene, which encodes the transcriptional regulator σ^54^ (also referred to as σ^L^). Similarly to other sigma factors, σ^54^ regulates genetic expression by incorporating the RNA polymerase complex and binding to specific promoter sequences, thus enabling selective transcription of a subset of genes (11–13). In the AA strain, *sigL* mutation leads to an amino acid substitution in a highly conserved DNA-binding region of the encoded protein. We hypothesize that this mutation results in the loss of nucleic acid binding capacity and thus impedes transcription of the σ^54^ regulon. According to *in silico* analyses, this regulon is apparently conserved in other *C. beijerinckii* strains. To confirm this hypothesis, *sigL* (Cbei_0595) was deleted in the acetone-butanol producer *C. beijerinckii* NCIMB 8052, and the resulting strain was shown to exhibit a solvent-deficient phenotype highly similar to the AA mutant obtained from the DSM 6423 strain. In addition to regulating solventogenesis, our experiments revealed that this sigma factor also controls utilization of alternative carbon sources, such as lactose and cellobiose, and is not required to complete the sporulation process, making it a central and specific controller of carbon metabolism in *C. beijerinckii*.

## Results

### The *C. beijerinckii* AA mutant metabolism is orientated towards acid production

Phenotypic characterization of a solventogenic *Clostridium* is usually assessed by the identification of fermentation products. We therefore performed triplicate fermentation assays of both *C. beijerinckii* DSM 6423 wild-type and AA strains to compare their product pattern after 48 hours in Gapes medium.

When compared to the wild-type strain, the AA mutant showed a very distinctive behavior. Firstly, growth was strongly impacted with a ca. 2-fold decrease in OD_600_ units after 48 hours of fermentation in Gapes medium, influencing end-products concentrations and glucose consumption (Fig. 1, Supplementary file S1).

**Figure 1:**
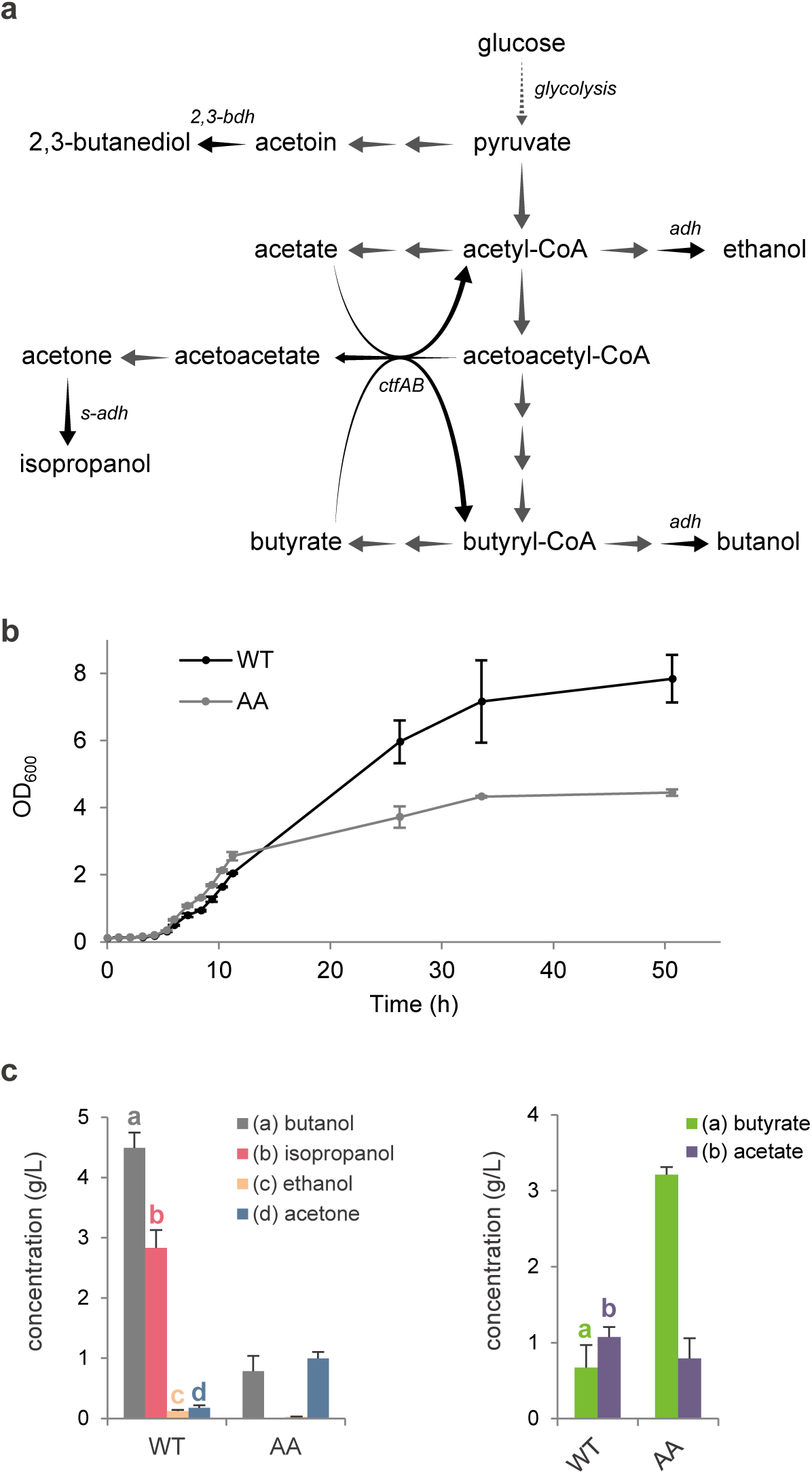
Comparative phenotypic analysis of *C. beijerinckii* DSM 6423 wild-type and AA strains. a. Simplified central metabolism of *C. beijerinckii* DSM 6423. b. Growth kinetics comparison between *C. beijerinckii* DSM 6423 (WT) and the AA mutant. c. Final solvent and acid concentrations measured after 48h of fermentation in Gapes medium for wild-type *C. beijerinckii* DSM 6423 and the AA mutant. Error bars indicate the standard deviation of triplicate experiments.

Although butanol was still produced, the central metabolism of the AA strain shifted towards acid formation (butyrate, acetate), with a final solvent:acid ratio of 1:3, compared to 6:1 in the wild-type strain (Fig. 1c). Acetate and butyrate are normally produced during the acidogenic phase and are further consumed during solventogenesis (14). In the AA mutant, increased acids production results in a substantial drop of the pH which may prematurely cause cell growth to abort. Since *C. beijerinckii* DSM 6423 has multiple copies of butanol and butyraldehyde dehydrogenases encoding genes in its genome, partial loss of butanol production capacity could be linked to the inactivation of one or several of these genes.

Isopropanol production was abolished in the AA strain but its precursor - acetone - was still detected. This tends to indicate that the expression of the sole secondary alcohol dehydrogenase (s-ADH) which notably catalyzes the acetone-to-isopropanol reaction is also affected in this mutant. Being linked to acid consumption, acetone formation suggests that butyrate and acetate uptake still occurs. In the wild-type DSM 6423 strain, the resulting acetyl-CoA and butyryl-CoA are further reduced into acetaldehyde/butyraldehyde and ethanol/butanol. Alcohol synthesis being drastically reduced in the AA strain, it may imply that most of the acetyl-CoA and butyryl-CoA are converted back to acetate and butyrate. This assumption would be supported by the increased (acetone+isopropanol):butanol ratio in the AA strain (close to 1:1, versus 1:3 in the wild-type). Indeed, this ratio is expected to be higher if butyryl-CoA is preferentially used to synthesize butyrate instead of butanol.

In summary, the AA strain mainly produces organic acids (butyrate, acetate), which probably results from the coupled inactivation of one or several gene(s) involved in isopropanol and butanol production.

### The AA mutant is still able to complete the sporulation process

Complete or partial loss of solventogenesis in the *C. beijerinckii* species is often associated with the degeneration phenomenon (i.e. loss of the capacity to form spores in addition to the loss of solvent production) (15). Given its phenotype, we therefore decided to investigate the sporulation ability of the AA mutant.

A comparative sporulation assay confirmed that sporulation ability was conserved, despite an approximate 10/100 fold decrease in the number of spores when compared to the wild-type (Supplementary file S1). However, given the limited growth of the AA strain, this decrease may not necessarily be linked to a genetic deficiency but could instead result from the shorter lifespan of the bacteria.

Importantly, demonstration of the sporulation ability in the AA strain makes its phenotype unrelated to the degeneration phenomenon, which suggests that its genome contains solventogenic-specific mutations.

### A punctual mutation in the *sigL* gene is predicted to impede solvent formation and sugar consumption

We next adopted a forward genetic approach to identify the gene(s) responsible for the AA phenotype. Whole-genome sequencing followed by read mapping on the wild-type genome allowed the identification of 13 SNPs (Table 2) along the chromosome of the strain. Among the affected genes, CIBE_0767 particularly retained our attention because BLAST analysis of its product against Swissprot database (using MaGe platform (16)) revealed a high similarity with a putative RNA polymerase σ^54^ factor (encoded by the *sigL* gene) of *Clostridium kluyveri* (17). In the AA strain, mutation of this gene causes a serine to phenylalanine substitution at position 366. The σ^54^ transcriptional factor has already been well described in several bacterial species and contains three major domains (12) (Fig. 2a). Region I (RI) is involved in core RNA polymerase, enhancer, and DNA binding. Region II (RII) is acidic and poorly conserved. Region III (RIII) is divided in several conserved sub-regions interacting with core RNA polymerase (CBD) and DNA at the −12 consensus motif (HTH) and the −24 motif (RpoN). *In silico* analysis of the *sigL* mutation detected in the AA mutant revealed that the amino acid substitution (S366F) is localized in the HTH motif, at an extremely conserved position (Fig. 2b). Given the significant difference in terms of steric hindrance, polarity and hydrophobicity resulting from a phenylalanine to serine substitution, this mutation may drastically impede the −12 motif recognition, yielding a partial or total inactivation of σ^54^.

**Table 2:** Mutations detected in the AA strain.

| Locus tag | Description | Change | Genome position | CDS Position | Amino Acid Change | Protein Effect |
| --- | --- | --- | --- | --- | --- | --- |
| CIBE_0366 | conserved protein of unknown function | C -> T | 390,092 | 2,215 | Q -> X | Truncation |
| CIBE_0767 | putative RNA polymerase sigma-54 factor | C -> T | 770,679 | 1,097 | S -> F | Substitution |
| CIBE_1279 | conserved exported protein of unknown function | C -> T | 1,347,126 | 31 | none | None |
| CIBE_1303 | conserved protein of unknown function | A -> G | 1,373,758 | 218 | Y -> C | Substitution |
| N/A<sup>a</sup> | intergenic | G -> A | 1,691,364 | N/A<sup>a</sup> | N/A<sup>a</sup> | N/A<sup>a</sup> |
| CIBE_2357 | Binding-protein-dependent transport systems inner membrane component | G -> A | 2,397,512 | 614 | G -> D | Substitution |
| CIBE_2358 | NMT1/THI5-like protein | G -> A | 2,397,959 | 138 | None | None |
| CIBE_2486 | protein of unknown function | C -> T | 2,535,980 | 103 | P -> S | Substitution |
| CIBE_2488 | protein of unknown function | C -> T | 2,537,804 | 157 | P -> S | Substitution |
| CIBE_3408 | putative hydrolase subunit antagonist of Kipl | C -> T | 3,476,677 | 206 | T -> I | Substitution |
| CIBE_3475 | Histidine kinase | C -> A | 3,572,491 | 732 | R -> S | Substitution |
| CIBE_4586 | dihydroxy-acid dehydratase | G -> T | 4,675,905 | 1,159 | V -> F | Substitution |
| N/A<sup>a</sup> | intergenic | C -> T | 4,931,169 | N/A<sup>a</sup> | N/A<sup>a</sup> | N/A<sup>a</sup> |
| CIBE_5453 | short chain acyl-CoA transferase: fused alpha subunit ; beta subunit | A -> G | 5,622,646 | 919 | S -> P | Substitution |
| CIBE_5584 | putative ABC transporter (ATP-binding protein) | G -> A | 5,774,626 | 146 | R -> K | Substitution |

<sup>a</sup> not applicable

**Figure 2:**
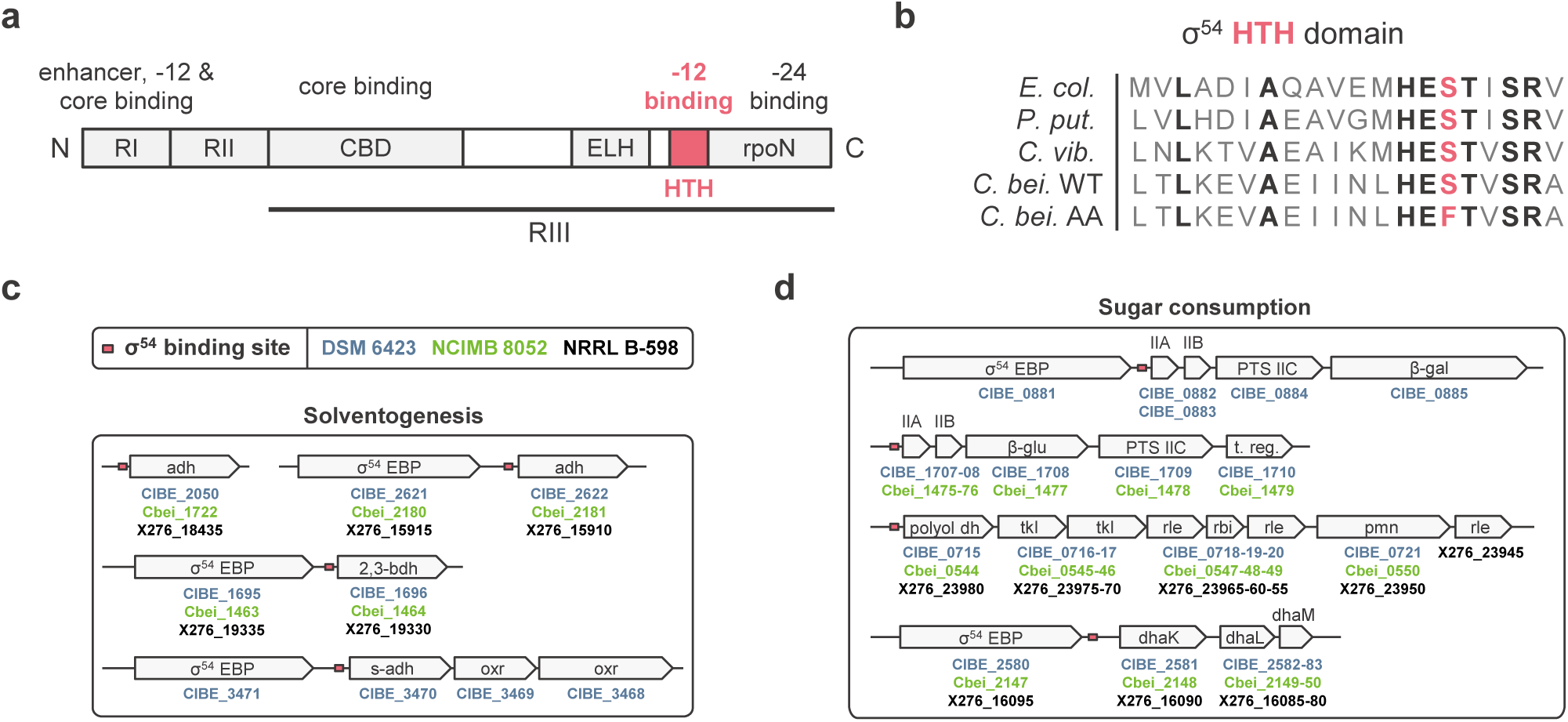
Structural position of σ^54^ mutation in *C. beijerinckii* DSM 6423 AA strain and predictions for this sigma factor role in several model strains. a. Structural organization of the σ^54^ sigma factor. RI-III: region I to III. CBD: core binding domain. ELH: extra-long α-helix. HTH: helix turn helix motif. rpoN: RNA polymerase factor N domain. b. Localization of the S336F mutation identified in the AA strain. *E. col*.: *Escherichia coli*; *P. put.*: *Pseudomonas putida*; *C. vib.*: *Caulobacter vibrioides*; *C. bei.*: *Clostridium beijerinckii*. c,d, *In silico* predictions of (c) solvent production and (d) sugar metabolism regulation by σ^54^ in several *C. beijerinckii* strains. Transcriptional units involved in these pathway bear an upstream σ^54^ binding site (consensus: TGGCANNNNNNTTGCW). adh: alcohol dehydrogenase; 2,3-bdh: 2,3-butanediol dehydrogenase; s-adh: secondary alcool dehydrogenase; oxr: oxidoreductase. PTS: phosphotransferase system; β-gal: β-galactosidase; β-glu: β-glucosidase; t. reg.: transcriptional regulator; dhaK/L/M: dihydroxyacetone kinase subunits K/L/M; polyol dh: polyol dehydrogenase; tkl: transketolase; rle: ribulose phosphate epimerase; rbi: ribose phosphate isomerase; pmn: phosphomannomutase.

We subsequently looked at the predicted σ^54^–regulon along the chromosome of *C. beijerinckii* DSM 6423. The σ^54^ binding motif consensus (TGGCANNNNNNTTGCW, based on the study of validated σ^54^ promoters by Barrios and colleagues (13)) was found at 57 genomic sites, 35 of which were localized up to a few hundred base pairs upstream of a coding sequence (Supplementary file S2). In particular, two genes coding for strongly expressed (8) alcohol dehydrogenases (CIBE_2050 and CIBE_2622, which catalyze the conversion of acetaldehyde and butyraldehyde into ethanol and butanol (18, 19)) as well as the secondary alcohol dehydrogenase s-ADH encoding gene (CIBE_3470, (18, 20, 21)) were identified (Fig. 2c). Although 2,3-butanediol is usually not observed in laboratory conditions, the only gene of the genome encoding a 2,3- butanediol dehydrogenase (CIBE_1696, similar to the one described by Raedts et al. (22)) was identified as being part of the regulon.

Unlike other sigma factors, σ^54^-bound RNA polymerase requires the ATPase activity of an adjacent Enhancer Binding Protein (EBP) to start transcription initiation (23, 24). Importantly, genes encoding σ^54^ EBPs were found in the direct vicinity of CIBE_2622, CIBE_3470 and CIBE_1696, which strengthens the hypothesis of a σ^54^-driven transcriptional control of their expression.

In order to investigate the functionality of the σ^54^-regulon, we used the synteny tool from MaGe platform (16) to compare the position of σ^54^ binding sites in *C. beijerinckii* DSM 6423 with two other *C. beijerinckii* model strains i.e. NCIMB 8052 and the recently reannotated NRRL B-598 (25, 26). Apart from the s-ADH encoding gene which drives isopropanol production, these putative σ^54^–dependent transcriptional units appeared to be conserved in these acetone-butanol producing strains. Besides, downregulation of the butanol and secondary alcohol dehydrogenases linked to σ^54^ potential inactivation would match the AA phenotype, which gives another meaningful hint that *sigL* mutation may be significant.

Interestingly, the rest of the predicted σ^54^-regulon mainly contains genes associated with control sugar uptake (mainly PTS-based transporters, for a review in solventogenic *Clostridia*, see Mitchell, 2015 (27)) and metabolism (Fig. 2d, Supplementary file S2). Lactose and cellobiose transport and metabolism are predicted to be regulated by σ^54^ in similar operons encoding a phosphotransferase system (PTS) for sugar transport and a hydrolase (β-galactosidase CIBE_0885; β-glucosidase CIBE_1708). Interestingly, a highly expressed and conserved operon encoding a dihydroxyacetone phosphate kinase complex (*dhaKLM*; CIBE_2581-2582-2583) also appears to be controlled by σ^54^. This enzyme can be involved in glycerol metabolism, or directly in the central carbon pathway when associated to 6P-fructose aldolases (28, 29). Notably, two 6P-fructose aldolases (CIBE_0334 and CIBE_0411), one of which being well expressed in the available RNA-seq dataset (8), are annotated in the DSM 6423 genome. Lastly, parts of the pentose phosphate pathway also seem to be regulated by σ^54^, with two conserved similar operons encoding transketolases, ribose/ribulose phosphate epimerases and isomerases (Fig. 2d and supplementary file S2).

In summary, analysis of the AA genotype revealed an amino acid substitution potentially inactivating a protein predicted not only to regulate solventogenesis but also other central metabolic pathway such as sugar consumption in several *C. beijerinckii* strains.

### Complementation with wild-type *sigL* gene fully restores solventogenesis and sugar metabolism in the AA strain

As several SNPs were detected in different genes in the AA strain, specific complementation assays were required to evaluate the importance of each mutation in the phenotype of the strain. Considering the predicted crucial role of the putative σ^54^-encoding gene (*sigL*, CIBE_0767), we decided to clone the wild-type coding sequence of *sigL* under the control of its endogenous promoter in the pFW01 vector, yielding the pFW-σ54 plasmid and to introduce it into both wild-type and AA strains. Fermentation analysis revealed that the recombinant AA pFW-σ54 strain displayed a wild-type solventogenic metabolism (Fig. 3). Indeed, normal acid and solvent production (Fig. 3a), as well as glucose consumption and biomass production (Supplementary file S3) were fully restored when compared to the wild-type strain containing the empty vector. These results confirmed that CIBE_0767, encoding the σ^L^ transcriptional factor, is drastically influencing the regulation of the central metabolism of the AA strain in our conditions. We therefore kept on investigating its impact on other predicted metabolic pathways.

**Figure 3:**
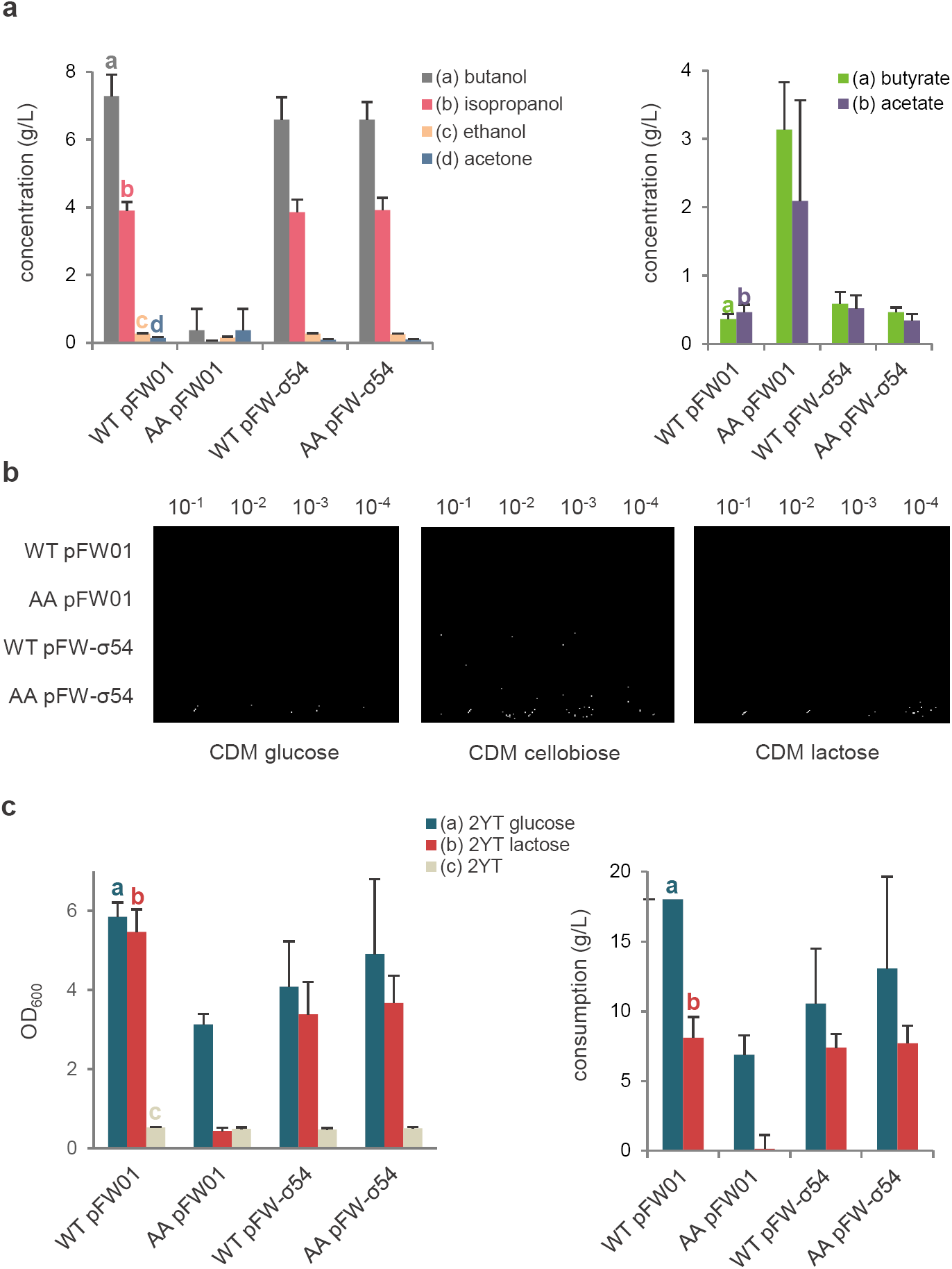
Phenotypic analysis of *C. beijerinckii* DSM 6423 σ54 complementation in the AA strain. a. Fermentation products after 72 hours in Gapes 60 g/L glucose medium of *C. beijerinckii* DSM 6423 wild-type (WT), AA strains containing an empty plasmid (pFW01) or the complementation plasmid (pFW01-σ54). b. Sugar utilization assay on CDM plates containing 20 g/L glucose, cellobiose or lactose. Pictures were taken after 24h of incubation on glucose and cellobiose, and 48h on lactose. c. Sugar utilization assay in liquid 2YT with 20 g/L glucose (2YTG), lactose (2YTL) or without added carbon source (2YT). Biomass (given as a measure of optical density at 600 nm, OD_600_) and carbon consumption are measured 24h after the beginning of the fermentation. Error bars indicate the standard deviation of triplicate experiments.

Plate assays on minimal medium were used to analyze carbon sources usage. In our hands, no effect after 24 or 48 hours was seen on bacterial growth on fructose, mannose, xylose, arabinose, sucrose, glycerol, inulin, starch or mannitol (data not shown). In contrast, growth was impeded with cellobiose and lactose in the AA control strain (empty vector), and restored when complemented with wild-type σ^54^ (Fig. 3b). However, quantification of this inhibition in liquid medium revealed unexpected results. Growth was not altered in liquid 2YT 20 g/L cellobiose (data not shown) whereas lactose did exert a strong effect on bacterial growth. After 24h of fermentation the AA pFW01 strain had grown similarly on 2YT 20 g/L lactose than on the 2YT control condition containing no supplementary carbon source (Fig. 3c). HPLC analysis confirmed that almost no lactose was consumed by the AA strain in this time frame. On the other hand, a normal growth on lactose was observed with the complementation strain with a consumption of ca. 8 g/L of lactose in 24h. The fermentation was carried out for 72 more hours, and end-point analysis revealed that lactose uptake inhibition was only partial (Supplementary file S4).

Thus, by complementing the AA strain with the wild-type version of the *sigL* gene, we demonstrated that the S366F point mutation in the σ^54^ is likely the unique modification in the AA strain that impacts solvent production and sugar assimilation. Besides, our results suggest that σ^54^ is a master regulator of these metabolic pathways in *C. beijerinckii* DSM 6423.

### Overexpression of the acid-reassimilation pathway does not restore a wild-type phenotype in the AA strain

To determine whether acid uptake might be impaired in the AA strain, we designed an experiment in which the acid reassimilation pathway would be overexpressed in the AA mutant. In order to do this, a pFW-FC06 plasmid was constructed to overexpress the *s-adh, ctfA*, and *ctfB* genes in an operonic structure. This operon was previously used in an overexpression plasmid (30) or integrated in the chromosome (31) to convert the model acetone-butanol producing organism *C. acetobutylicum* ATCC 824 into a performant isopropanol-butanol producer.

Introduction of the pFW-FC06 vector slightly increased solvent production and glucose consumption in the AA strain, compared to the introduction of an empty vector (supplementary file S5). However, even if acetone was fully converted to isopropanol, the recombinant AA strain containing pFW-FC06 still accumulated butyrate and acetate at levels comparable to the AA mutant. The latter observation suggests that, consistently with *in silico* predictions, the CtfA/B complex is still functional in the AA mutant (i.e. overexpression of *ctfA/B* does not have much effect on acid levels, similarly to the wild-type strain). The main metabolism roadblock thus probably comes from the aforementioned deficient butanol synthesis pathway. Indeed, in the AA strain, this result would be expected if cyclic conversion of butyryl-CoA and acetyl-CoA to butyrate and acetate, and back to acyl-CoAs via the CtfA/B enzyme, was occurring.

In summary, this experiment gives an additional insight on the molecular mechanisms triggering solventogenesis in *C. beijerinckii*: by regulating alcohol dehydrogenase expression, σ^54^-mediated transcription might be able to modulate the balance between acidogenesis and solventogenesis.

### CRISPR/Cas9-mediated deletion of *sigL* in *C. beijerinckii* NCIMB 8052 results in an AA-like phenotype

Complementation with wild-type σ^54^ in the AA strain shed light on *sigL* role in solventogenesis and sugar uptake. However, given the many mutations found along the genome of the AA mutant, we aimed at confirming the effect of *sigL* inactivation by deleting it in a wild-type strain. This approach also provides a convenient way to compare the effects of the S336F punctual mutation to a complete gene deletion. The model strain *C. beijerinckii* NCIMB 8052 was chosen as a chassis for this modification since, unlike the DSM 6423 strain, markerless genome editing techniques (in particular CRISPR-derived tools) have already been demonstrated to be effective in this microorganism (32–34).

We thus designed a CRISPR/Cas9 approach to inactivate *sigL* in the NCIMB 8052 strain. An inducible system based on anhydrotetracycline (aTc) addition was used, based on the pCas9_ind_ vector previously used in *C. acetobutylicum* ATCC 824 (31). Homology regions for genome editing were designed to delete most of the *sigL* gene, resulting in a severely truncated protein (13 amino acid residues, versus 463 for the full gene; Fig. 4a). Following insertion of those homology sequences and of an anhydrotetracycline inducible gRNA cassette, the resulting pCas9-Δ*sigL* plasmid was introduced into *C. beijerinckii* NCIMB 8052 by electroporation. Transformants were exposed to anhydrotetracycline, which allowed the selection of a Δ*sigL* mutant (Fig. 4b). This strain was subsequently complemented with the pFW-σ54 and pFW-σ54-AA plasmids, the latter allowing the expression of a S336F-mutated σ^54^.

**Figure 4:**
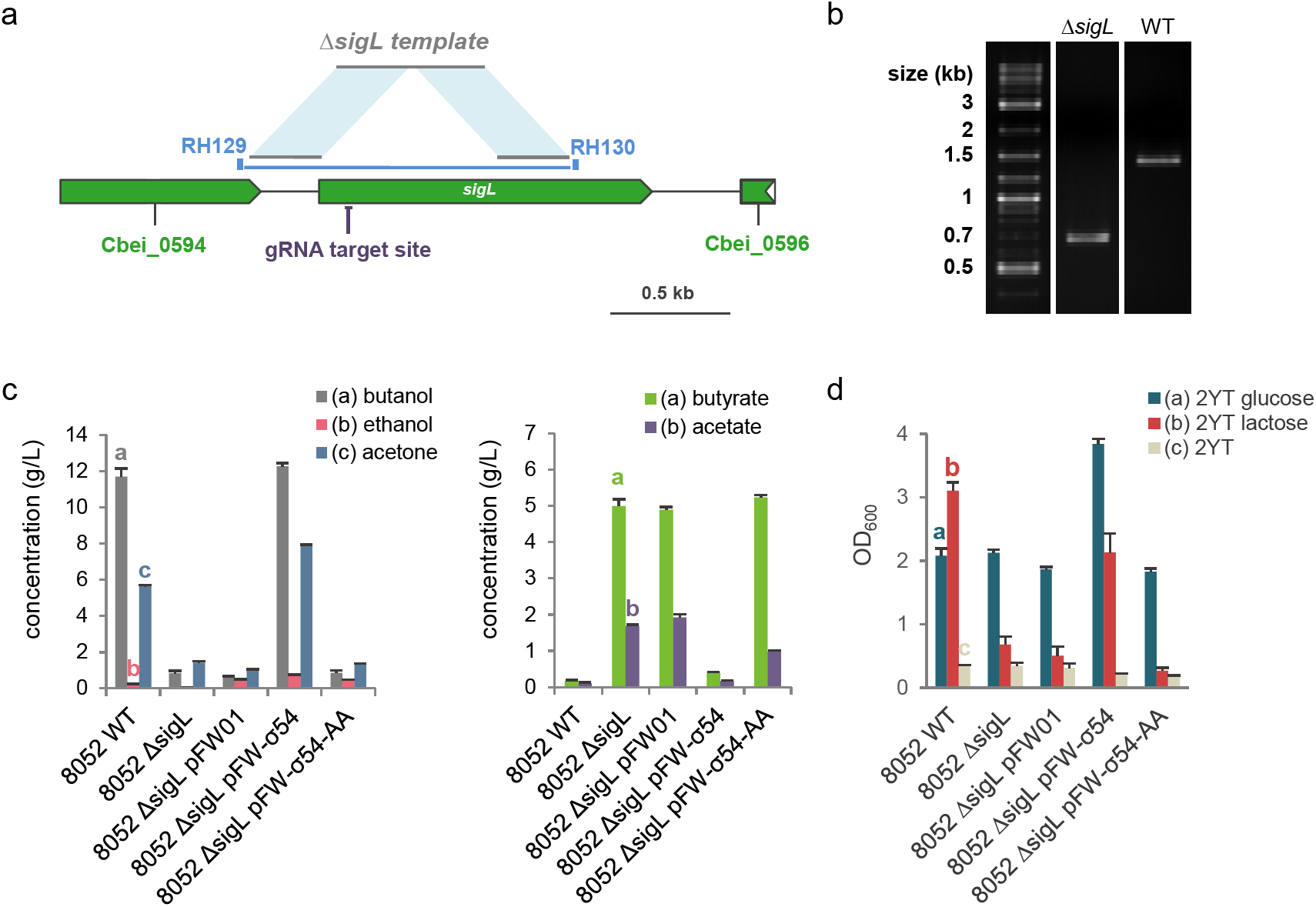
Design of CRISPR/Cas9-meditated σ^54^ deletion in *C. beijerinckii* NCIMB 8052 and phenotypic comparison of wild-type and mutant σ^54^ complementation. a. *sigL* (Cbei_0595) genomic region with CRISPR/Cas9 genome editing design. b. Verification by colony PCR of *sigL* deletion with primers RH129 and RH130 encompassing the deleted region. Expected band sizes are 1406 (wild-type) and 670 (Δ*sigL*) bp. c. Comparative analysis of fermentation capacities of *C. beijerinckii* NCIMB 8052 wild-type, Δ*sigL* and complemented strains. d. Growth comparison of *C. beijerinckii* NCIMB 8052 wild-type, Δ*sigL* and complemented strains after 24 hours of fermentation in 2YT, 2YTG and 2YTL liquid media. gRNA: guide RNA. Error bars indicate the standard deviation of triplicate experiments.

Fermentation assays revealed that solvent production was drastically altered in the Δ*sigL* strain, and that a wild-type phenotype could only be restored with pFW-σ54 (Fig. 4c, Supplementary file S7). Complementation with pFW-σ54-AA notably appeared to have no effect on acid reassimilation and solvent production when compared to the Δ*sigL* empty vector control strain. Besides, all of those strains displayed a resistance to heat shock (cells scraped from one-week-old plates, 80°C for 10 min, data not shown), highlighting their retained capacity to correctly form spores. Lastly, Δ*sigL* mutant growth appeared to be highly inhibited in lactose-based medium, similarly to the AA strain (Fig. 4d). Conversely, cellobiose uptake did not seem to be impacted in this strain (data not shown).

In addition to confirming our hypothesis on the role of σ^54^ for solventogenesis and sugar utilization, deleting *sigL* in the NCIMB 8052 model strain allowed us to expand our conclusions regarding this transcriptional factor to other microorganisms belonging to the *C. beijerinckii* species.

## Discussion

In this study, the allyl alcohol mutant generated from *C. beijerinckii* DSM 6423 by Máté de Gerando et al. (9) was genetically characterized by a forward genetic approach. Complementation assays demonstrated that the AA phenotype is due to a unique point mutation in the CIBE_0767 *sigL* gene encoding the σ^54^ transcriptional factor. This mutation leads to the substitution of a highly conserved serine in the HTH domain by a phenylalanine residue (S336F). Given the central role of the HTH domain in DNA recognition (35, 36), this mutation is likely to strongly impede the σ^54^-RNA polymerase complex binding to promoter sequences and thus to inhibit transcription of the σ^54^-regulon. If the effect of this particular mutation on σ^54^ functionality has not been described so far, Coppard and Merrick (36) performed targeted mutagenesis of the corresponding serine of the HTH domain of *K. pneumoniae*. Its substitution by most residues (especially large ones like lysine or tryptophane) completely shut down σ^54^-mediated transcription, suggesting this serine is crucial for a functional activity, which supports our hypothesis that σ^54^ is inactivated in the AA strain.

The deleterious effects of allyl alcohol on alcohol-producing microorganisms (yeast (37–39), *E. coli* (40)) and its tendency to generate spontaneous mutations have been reported decades ago by early workers. Reverse activity of alcohol dehydrogenases catalyzes the formation of the highly toxic acrolein molecule from allyl alcohol, enabling the selection of alcohol dehydrogenase deficient mutants. In *Clostridia*, similar work was pioneered by Dürre and colleagues (10), who obtained *C. acetobutylicum* mutants unable to produce butanol. Interestingly, butyraldehyde dehydrogenase activity was drastically reduced in these strains. Given the *in silico* σ^54^-targeted promoter predictions, σ^54^ apparently does not control the butyraldehyde dehydrogenase genes transcription in the DSM 6423 strain. The supposed inactivation of σ^54^ in the AA mutant rather seems to lead to a decreased expression of two major alcohol dehydrogenases genes, resulting in a phenotype close to what was observed in the *C. acetobutylicum* corresponding mutants. Alcohol dehydrogenase expression being driven by σ^54^, inactivation of *sigL* would result in an accumulation of acids, whose CoA-derivatives obtained via the CoA-transferase CtfA/B would be preferentially transformed back into acetate and butyrate via the phosphotransbutyrylase/acetylase – butyrate/acetate kinase pathway. Accumulation of acids in the fermentation broth could also be linked to a defect in CoA-transferase activity. However, overexpression of the acid reassimilation pathway (CtfA/B, s-ADH) in the AA strain did not increase acid uptake. In accordance with *in silico* predictions, this experiment suggests σ^54^ mainly drives the regulation of solventogenesis by modulating alcohol dehydrogenase activity.

σ^54^ was also shown to control sugar uptake and usage, consistently with what Nie et al. predicted for the *Clostridium* genus (41). In particular, growth was limited on cellobiose and lactose in the AA strain, though cellobiose inhibition could only be observed on minimal medium. This may be explained by the existence of a duplicate cellobiose assimilation pathway in the DSM 6423 genome. Different copies of these genes might be 1) not controlled by σ^54^ and/or 2) poorly expressed on solid minimal medium. The presence of two alternative σ^54^-independent β-glucosidase genes (CIBE_5551 and CIBE_5773) in the *C. beijerinckii* DSM 6423 genome may support this assumption.

The impact of the S336F mutation in the σ^54^ sequence was later investigated by introducing expression plasmids of either the wild-type or the S336F version of σ^54^ in a Δ*sigL* mutant of the model strain *C. beijerinckii* NCIMB 8052. The Δ*sigL* mutant displayed a phenotype highly similar to the AA strain. Indeed, solventogenesis was severely impaired which resulted in an acid - more particularly butyrate - orientated metabolism. Importantly, complementation with pFW-σ54 fully restored solventogenesis, while pFW-σ54-AA had no impact on the strain ability to reduce its acids. Besides, *sigL* deletion was shown to severely impact lactose uptake, similarly to the AA mutant. These experiments overall demonstrated that the S336F mutation completely inactivates σ^54^ and are thus consistent with our previous results.

σ^54^ belongs to a unique category of sigma factors, being evolutionary distinct to the σ^70^ family, which comprises all of the other sigma factors (42, 43). This difference implies a very different mode of action for the σ^54^-RNA polymerase complex: unlike its σ^70^ counterpart, the holoenzyme is unable to initiate transcription without the help of an enhancer binding protein (EBP) (24). EBPs are often acting several hundred of base pairs upstream of the σ^54^-holoenzyme deposition sites and contain a central AAA+ domain. The latter provides, through ATP hydrolysis, the energy necessary for structural rearrangements within the σ^54^-bound holoenzyme, which in turn allows transcription initiation. EBP control on σ^54^-dependant transcription is mainly exerted by a signal-sensing domain, which upon specific environmental stimuli activates or represses its ATPase activity (24). This enhancer-dependent system offers a very tight control on gene expression, and σ^54^ regulons are often associated with various biological processes that require a stringent control (e.g. virulence (44–46) or biofilm formation (47–49)). Multiple σ^54^ EBPs were found in *C. beijerinckii* genomes and are thus likely to be the primary effectors of σ^54^-mediated regulation. Indeed, transcript abundance analysis (8) shows that *sigL* is not differentially expressed over the time-course of a batch fermentation, whereas its target solventogenic genes (CIBE_2050, CIBE_2622, CIBE_3470) are among the top upregulated genes during the transition from acidogenesis to solventogenesis, which suggests that activation of their transcription is mediated by other factors (i.e. the enhancer binding proteins). A well-known example of such regulation is the σ^54^-driven transcription of the levanase operon in *B. subtilis*, which encodes an enzymatic and transport machinery allowing the degradation and consumption of fructose polymers (50, 51). The enhancer binding protein LevR was shown to mediate the activation and repression of the expression of this operon by a complex interplay with the PTS system (54). However, in *C. beijerinckii*, the exact stimuli recognized by these effectors and underlying regulatory mechanisms that allow transcription of σ^54^ target genes still remain to be clarified.

In bacteria, the housekeeping sigma factor (e.g. σ^70^ in *E. coli*) integrates RNA polymerase and drives transcription of most housekeeping genes. However, bacterial genomes encode multiple alternative sigma factors which direct RNA polymerase to different promoters, providing a simple but broadly exploited strategy to regulate genetic expression (52, 53). In solventogenic *Clostridia*, alternative sigma factors have mainly been highlighted for their crucial role in the regulation of the multi-stages sporulation cascade (55–57). Inactivation of the sporulation regulating factors σ^E^ and σ^F^ was shown to block spore formation but also to impair solventogenesis in *C. acetobutylicum* when cultures of σ^E^ and σ^F^ knockout strains were inoculated with mid-exponential phase precultures. Using stationary phase precultures, this phenomenon was not observed. On the other hand, σ^K^ plays in *C. acetobutylicum* an early role in the sporulation process by controlling the expression of the solventogenesis / sporulation major regulator Spo0A, and its inactivation thus leads to the loss of solvent production and spore formation abilities. These striking examples of sigma factor mediated regulation underline the intricate relationship between solventogenesis and sporulation. In the ballet of transcriptional and post-transcriptional factors regulating their outcome (58), the discovery of the role of another peculiar sigma factor controlling solventogenesis is thus not surprising. Indeed, the absolute requirement of an enhancer binding protein (24) makes σ^54^ stand out for its efficiency to tightly control transcription of highly regulated metabolic pathways such as solventogenesis. Interestingly, no sporulation-specific targets could be found in the predicted σ^54^ regulon, and σ^54^ inactivation did not drastically inhibit the sporulation cascade, which makes it a carbon-specific regulator.

In summary, we described σ^54^ as a master regulator of solventogenic pathways in the isopropanol/butanol producer *C. beijerinckii* DSM 6423 and this function was predicted to be conserved in other model strains of the *C. beijerinckii* species. This transcriptional factor was also found to control sugar consumption and is therefore an essential controller of carbon metabolism in *C. beijerinckii*.

## Material & Methods

### Strains, media and culture conditions

Strains and plasmids used in this study are presented in Table 1.

**Table 1:**
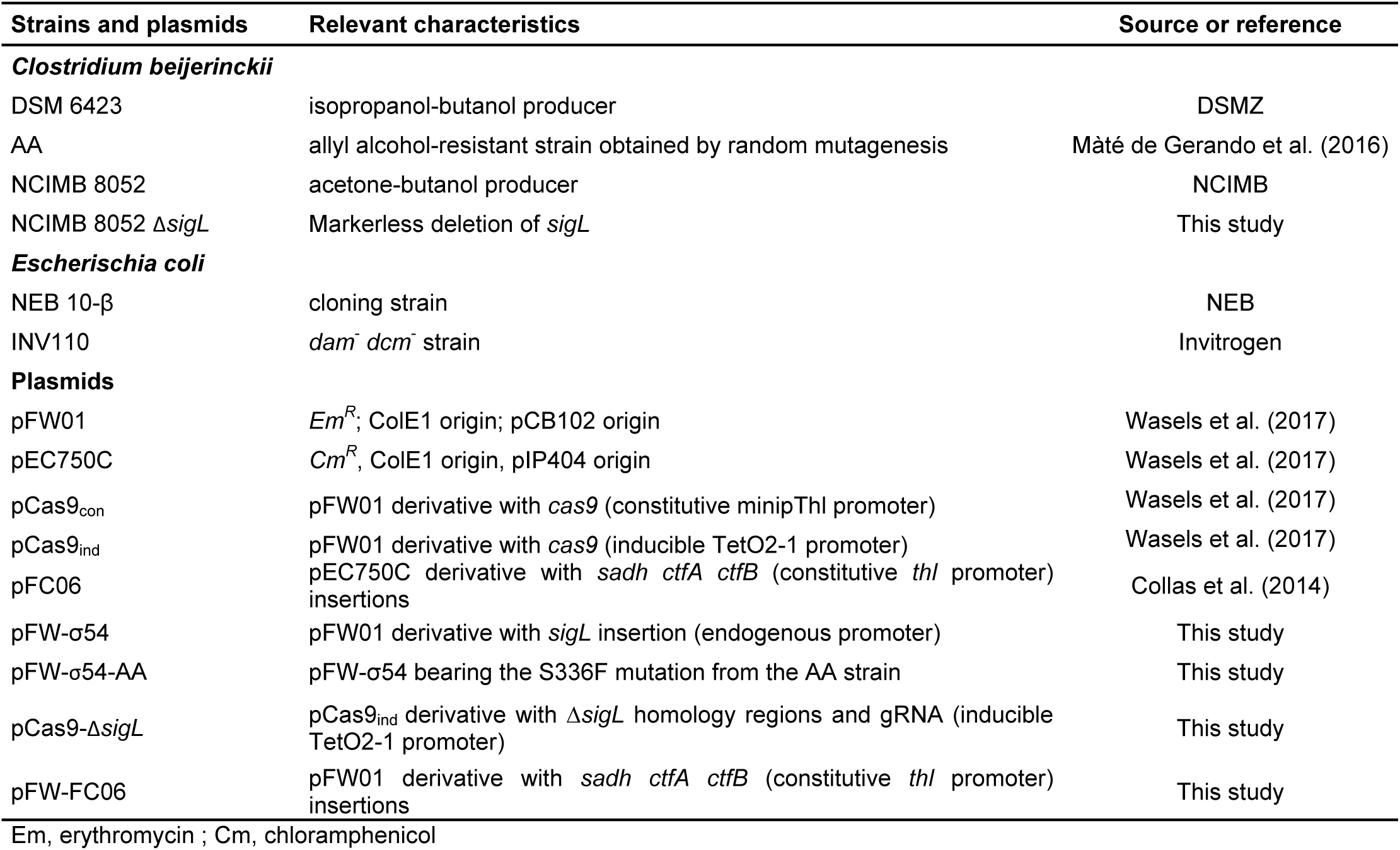
Strains and plasmids used in this study.

| Strains and plasmids | Relevant characteristics | Source or reference |
| --- | --- | --- |
| ***Clostridium beijerinckii*** |  |  |
| DSM 6423 | isopropanol-butanol producer | DSMZ |
| AA | allyl alcohol-resistant strain obtained by random mutagenesis | Mâté de Gerando et al. (2016) |
| NCIMB 8052 | acetone-butanol producer | NCIMB |
| NCIMB 8052 $\Delta sigL$ | Markerless deletion of *sigL* | This study |
| ***Escherichia coli*** |  |  |
| NEB 10-β | cloning strain | NEB |
| INV110 | *dam*<sup>-</sup> *dcm*<sup>-</sup> strain | Invitrogen |
| **Plasmids** |  |  |
| pFW01 | *Em*<sup>R</sup>; ColE1 origin; pCB102 origin | Wasels et al. (2017) |
| pEC750C | *Cm*<sup>R</sup>, ColE1 origin, pIP404 origin | Wasels et al. (2017) |
| pCas9<sub>con</sub> | pFW01 derivative with *cas9* (constitutive minipThl promoter) | Wasels et al. (2017) |
| pCas9<sub>ind</sub> | pFW01 derivative with *cas9* (inducible TetO2-1 promoter) | Wasels et al. (2017) |
| pFC06 | pEC750C derivative with *sadh ctfA ctfB* (constitutive *thl* promoter) insertions | Collas et al. (2014) |
| pFW-σ54 | pFW01 derivative with *sigL* insertion (endogenous promoter) | This study |
| pFW-σ54-AA | pFW-σ54 bearing the S336F mutation from the AA strain | This study |
| pCas9-$\Delta sigL$ | pCas9<sub>ind</sub> derivative with $\Delta sigL$ homology regions and gRNA (inducible TetO2-1 promoter) | This study |
| pFW-FC06 | pFW01 derivative with *sadh ctfA ctfB* (constitutive *thl* promoter) insertions | This study |

Em, erythromycin ; Cm, chloramphenicol

*Clostridium beijerinckii* was grown in liquid 2YTG (per liter: 16 g tryptone, 10 g yeast extract, 5 g NaCl, 20 g glucose, pH 5.2). Solid media was prepared with 15 g/L agar and less glucose (5 g/L). Strains were cultivated in an anaerobic chamber (Bactron) at 34°C without shaking.

*Escherichia coli* was grown in liquid or solid LB media, in aerobic conditions at 37°C with 180 rpm agitation when necessary.

### Whole-genome sequencing

Genomic DNA of the AA strain was purified with the GenElute Bacterial Genomic DNA Kit (Sigma-Aldrich) and was subsequently sequenced on an Illumina MiSeq sequencer (2 x 250 paired-end reads). Mutations were detected by read mapping against the *C. beijerinckii* DSM 6423 genome (8) using Geneious R10 (59). Raw reads have been deposited to the SRA database with the accession number XX.

### Predictions of σ^54^-regulons

Genomes of the *C. beijerinckii* DSM 6423, NRRL B-598 and NCIMB 8052 strains were scanned for the σ^54^ consensus motif (TGGCANNNNNNTTGCW (13), no mismatches allowed) using Geneious R10 (59). Genes were considered potentially regulated by σ^54^ if motifs were found up to 500 bp upstream their coding sequences. Predicted candidate genes/operons were then compared in the three strains using the synteny tool from MaGe platform (16).

### Fermentation assays and analytical methods

For fermentation assays, modified Gapes medium (60) was used. This medium contains per liter: 2.5 g yeast extract, 1 g KH_2_PO_4_, 0.6 g K_2_HPO_4_, 1 g MgSO_4_, 7 H_2_0, 6.6 mg FeSO_4_, 7 H_2_0, 0.1 g para-aminobenzoic acid, 2.9 g ammonium acetate, 60 or 80 g glucose. The medium was supplemented with erythromycin (20 µg/mL) when appropriate.

Fermentations were performed in triplicate. For each biological replicate, several clones isolated in 2YTG agar plates were used to inoculate 5 mL Gapes medium. These precultures were carried out at 34°C overnight in the anaerobic chamber. Serum flasks containing 20 mL Gapes medium were then inoculated with 2 mL of preculture and sealed with rubber stoppers. A pressure relief valve system was punctured through the rubber stoppers to prevent overpressure, and the serum bottles were incubated 48-72 hours at 34°C with agitation outside of the anaerobic chamber.

After OD_600_ (UV-1800 spectrophotometer, Shimadzu) was measured, samples from the fermentation were centrifuged 5 min at 8000 g. The supernatant was diluted with an internal standard (1-propanol: final concentration 0.5 g/L). Metabolites concentrations in the supernatant were quantified by chromatography.

Gas chromatography (Porabond-Q column from Agilent technologies, 25 m length, 0.32 internal diameter, 5 µm film thickness coupled to a flame ionization detector) was performed to determine solvent concentrations. Helium was used as a carrier gas at a flowrate of 1.6 mL/min. Column was gradually heated from 50°C to 250°C in a 30 min run.

Acid concentrations were quantified by HPLC (Aminex HPX-87H from Biorad coupled to a Spectra System RI 150 refractometer and a Waters 2487 Dual λ UV detector set at 210 nm). 0.01 M sulfuric acid mobile phase was used at a flowrate of 0.6 mL/min. Column temperature was set at 60°C.

Residual sugar quantities were determined by HPLC using an Aminex HXP-87P (Bio-Rad) coupled to a Varian 350 RI refractometer for detection. Water was used as a mobile phase at a flowrate of 0.4 mL/min. When quantifying sugar concentration for 2YT-based samples, a Micro-Guard De-Ashing Refill cartridge (Bio-Rad) was added to the chromatographic system for prior desalting of the samples. Column temperature was set at 80°C.

### Plasmid construction and transformation

All primers used in this study are listed in supplementary file S6.

The *sigL* gene (CIBE_0767) from *C. beijerinckii* DSM 6423 together with its endogenous promoter were amplified by PCR with primers RH077 and RH078. The pFW01 backbone as well as the thiolase terminator were amplified by PCR from plasmid pCas9_con_ (31) with primers RH086 and RH087. Those two amplicons were subsequently assembled by Gibson assembly (NEB) (61) to yield pFW-σ54. From this plasmid, pFW-σ54-AA was obtained by site-directed mutagenesis with primers RH136 and RH137.

The *sadh* gene (CIBE_3470) from *C. beijerinckii* DSM 6423 and the *ctfA* and *ctfB* genes from *C. acetobutylicum* ATCC 824 were amplified by PCR with primers MKz01 and MKz02 from plasmid pFC006 (30). The resulting PCR product was cloned between the KpnI and SalI restriction sites to obtain the pFW-FC06 vector.

For the deletion of *sigL* in *C. beijerinckii* NCIMB 8052, two homology regions and an anhydrotetracycline-inducible gRNA expressing cassette were first synthesized by BaseClear in a pUC57 plasmid. Homology regions were designed to delete a 736 bp fragment in the *sigL* gene, yielding a truncated ORF (13 amino acid residues instead of 463). This cassette was amplified with primers RH125 and RH126 and subsequently cloned into the pCas9_ind_ vector (31) at the XhoI restriction site. The gRNA protospacer (designed using Genious R10 (59)) was then introduced in the resulting vector by Golden Gate assembly at the BsaI restriction site with primers guide_sigL_fwd and guide_sigL_rev, yielding pCas9-Δ*sigL*.

The plasmids were then isolated and electroporated into *C. beijerinckii* as previously described (62, 63). *dam*^+^ *dcm*^+^ DNA and *dam*^-^ *dcm*^-^ DNA were used for *C. beijerinckii* NCIMB 8052 and DSM 6423, respectively.

### CRISPR/Cas9 genome edition

Following transformation of the pCas9-Δ*sigL* plasmid, Δ*sigL* mutants were obtained similarly to what was described by Wasels et al (31). Briefly, transformants were resuspended in liquid 2YTG (pH 5.2, 20 g/L glucose) and serially diluted for spotting onto 2YTG plates containing erythromycin (20 µg/mL) and anhydrotetracycline (50 ng/mL). Isolated colonies were tested by colony PCR with primers RH129 and RH130, encompassing the homology regions in the genome (Fig. 4a). PCR was expected to yield products of 670 and 1406 bp for deletion and wild-type strains, respectively (Fig. 4b). Positive mutants were subsequently cured by streaking them twice on 2YTG plates supplemented with 50 ng/mL anhydrotetracycline. Plasmid presence in isolated colonies was next tested on 2YTG plates containing erythromycin (20 µg/mL), and erythromycin-sensitive clones were selected for further analysis.

### Sporulation assays

Sporulation assays were based on survival to heat shock, as similarly described (55). Briefly, triplicates precultures were carried out overnight in 2YTG medium. The following day, serum bottles containing 20 mL 2YTG (15 g/L glucose, pH 6.8) were inoculated with 2 mL of preculture and subsequently sealed. After 5 days at 34°C without agitation, 1 mL of culture was heat shocked (10 min, 80°C) and serially diluted. 5 µL spots were made for each dilution on 2YTG plates, which were then incubated overnight at 34°C.

### Sugar utilization assays

For liquid sugar utilization tests, liquid 2YTG (pH 6.8) was used, as well as 2YTL (pH 6.8, 20 g/L lactose) and 2YT (pH 6.8, without purified carbon source). Precultures were carried out overnight from fresh colonies in 2YTG medium (pH 6.8) containing 5 g/L glucose and 20 µg/mL erythromycin at several dilutions (up to 10^-4^ dilution factor). The following day, fresh precultures were used to inoculate 20 mL (AA strain; serum bottles) / 5 mL (Δ*sigL* strain; 24 deep-well plates) of 2YTG, 2YTL or 2YT media with or without (as needed) erythromycin 20 µg/mL at 0.01 units of OD_600_. Samples were taken after 24 and 96 hours, and sugar concentration was quantified as described above.

Minimal medium (CDM) was used to visualize sugar uptake inhibition by the AA strain. This medium, described by Vasconcelos et al. (64), was supplemented with 15 g/L agar and contained 20 g/L glucose, lactose or cellobiose. Single colonies picked on CDM glucose (5 g/L) plates were serially diluted in water. Spots were made with 5 µL of the 10^-1^, 10^-2^, 10^-3^ and 10^-4^ dilutions on minimal medium plates containing glucose, cellobiose or lactose (20 g/L). Plates were visualized after 24 h (glucose, cellobiose) or 48 h (lactose).

## Acknowledgements

We are grateful to Michelle Kuntz for constructing the pFW-FC06 plasmid and to Gwladys Chartier for technical assistance.

